# Promoter activity-based case-control association study on *SLC6A4* highlighting hypermethylation and altered amygdala volume in male patients with schizophrenia

**DOI:** 10.1101/2020.05.06.058792

**Authors:** Tempei Ikegame, Miki Bundo, Naohiro Okada, Yui Murata, Shinsuke Koike, Hiroko Sugawara, Takeo Saito, Masashi Ikeda, Keiho Owada, Masaki Fukunaga, Fumio Yamashita, Daisuke Koshiyama, Tatsunobu Natsubori, Norichika Iwashiro, Tatsuro Asai, Akane Yoshikawa, Fumichika Nishimura, Yoshiya Kawamura, Jun Ishigooka, Chihiro Kakiuchi, Tsukasa Sasaki, Osamu Abe, Ryota Hashimoto, Nakao Iwata, Hidenori Yamasue, Tadafumi Kato, Kiyoto Kasai, Kazuya Iwamoto

**Affiliations:** Department of Neuropsychiatry, Graduate School of Medicine, The University of Tokyo, Tokyo, Japan; Department of Molecular Brain Science, Graduate School of Medical Sciences, Kumamoto University, Kumamoto, Japan; PRESTO, Japan Science and Technology Agency, Tokyo, Japan; International Research Center for Neurointelligence (WPI-IRCN), The University of Tokyo Institutes for Advanced Study (UTIAS), The University of Tokyo, Tokyo, Japan; UTokyo Institute for Diversity and Adaptation of Human Mind (UTIDAHM), The University of Tokyo, Tokyo, Japan; Department of Neuropsychiatry, Graduate School of Medical Sciences, Kumamoto University, Kumamoto, Japan; Department of Psychiatry, Fujita Health University School of Medicine, Aichi, Japan; Department of Child Neuropsychiatry, Graduate School of Medicine, The University of Tokyo, Tokyo, Japan; Division of Cerebral Integration, National Institute for Physiological Sciences, Aichi, Japan; Division of Ultrahigh Field MRI, Institute for Biomedical Sciences, Iwate Medical University, Iwate, Japan; Schizophrenia Research Project, Tokyo Metropolitan Institute of Medical Science, Tokyo, Japan; Heartful Kawasaki Hospital, Kanagawa, Japan; Institute of CNS Pharmacology, Tokyo, Japan; Laboratory of Health Education, Graduate School of Education, The University of Tokyo, Tokyo, Japan; Department of Radiology, Graduate School of Medicine, The University of Tokyo, Tokyo, Japan; Department of Pathology of Mental Diseases, National Institute of Mental Health, National Center of Neurology and Psychiatry, Tokyo, Japan; Osaka University, Osaka, Japan; Department of Psychiatry, Hamamatsu University School of Medicine, Shizuoka, Japan; Laboratory for Molecular Dynamics of Mental Disorders, RIKEN CBS, Saitama, Japan

**Keywords:** DNA methylation, CpG island shore, serotonin transporter, 5-HTTLPR, brain imaging, major psychosis

## Abstract

Associations between altered DNA methylation of the serotonin transporter (5-HTT)-encoding gene *SLC6A4* and early life adversity, mood and anxiety disorders, and amygdala reactivity have been reported. However, few studies have examined epigenetic alterations of *SLC6A4* in schizophrenia (SZ). We examined CpG sites of *SLC6A4*, whose DNA methylation levels have been reported to be altered in bipolar disorder, using three independent cohorts of patients with SZ and age-matched controls. We found significant hypermethylation of a CpG site in *SLC6A4* in male patients with SZ in all three cohorts. We showed that chronic administration of risperidone did not affect the DNA methylation status at this CpG site using common marmosets, and that *in vitro* DNA methylation at this CpG site diminished the promoter activity of *SLC6A4*. We then genotyped the 5-HTT-linked polymorphic region (5-HTTLPR) and investigated the relationship among 5-HTTLPR, DNA methylation, and amygdala volume using brain imaging data. We found that patients harboring low-activity 5-HTTLPR alleles showed hypermethylation and they showed a negative correlation between DNA methylation levels and left amygdala volumes. These results suggest that hypermethylation of the CpG site in *SLC6A4* is involved in the pathophysiology of SZ, especially in male patients harboring low-activity 5-HTTLPR alleles.

## Introduction

The serotonin transporter (5-HTT), encoded by *SLC6A4*, is a major monoamine transporter that regulates serotonin (5-HT) neurotransmission at the synaptic cleft, affecting emotions and stress responses;^1^ thus, 5-HTT is a target protein of antidepressants. *SLC6A4* has a functional polymorphism at a promoter region known as the 5-HTT-linked polymorphic region (5-HTTLPR) that is classified into short (S) and long (L) alleles,^2^ with the S allele exhibiting weaker transcriptional activity.^3,4^ Caspi *et al*.^5^ reported that gene-by-environment (G x E) interactions between 5-HTTLPR and adverse life experiences are associated with the development of anxiety and depression. To date, numerous case-control association studies have been performed, and several meta-analyses have validated the G x E interactions.^6-9^ However, their existence remains controversial,^10,11^ and the largest meta-analysis^12^ as well as the largest case-control study^13^ failed to replicate the findings.

In addition to 5-HTTLPR, altered DNA methylation of the promoter region of this gene has been reported to be associated with childhood maltreatment,^14-17^ bullying,^18^ low socioeconomic status,^19,20^ stressful life events,^21,22^ suicide,^23^ depressive symptoms,^24-26^ major depression,^27-30^ and bipolar disorder (BD).^31^ Although patients with schizophrenia (SZ) frequently exhibit depressive symptoms in various stages of the illness, few studies have analyzed the DNA methylation status of *SLC6A4* in SZ.^32^

We previously identified hypermethylation of *SLC6A4* in the context of BD through promoter-wide screening using lymphoblastoid cell lines (LCLs) derived from monozygotic twins discordant for BD^31^. Hypermethylation was detected at two CpG sites (chr17:30,235,246-30,235,247 and chr17:30,235,271-30,235,272, named CpG3 and CpG4, respectively) within the CpG island shore (defined as the region within 2 kb of a CpG island) in the promoter region of *SLC6A4*. We confirmed that hypermethylation of the two CpG sites existed in LCLs and in postmortem prefrontal cortices from individuals with BD^31^.

In this study, we first confirmed the previous finding of hypermethylation of *SLC6A4* in BD using peripheral blood cells (PBCs). We then examined DNA methylation levels in SZ and found that CpG3 was also hypermethylated in PBCs from patients with SZ in three cohorts. Animal model experiments using common marmosets suggested that this epigenetic change was unlikely to be the result of medication. *In vitro* methylation analysis revealed that DNA methylation of CpG3 resulted in loss of promoter activity in the cultured cell lines. To examine the relationship between 5-HTTLPR and CpG3 DNA methylation, we genotyped 5-HTTLPR in detail. Strikingly, we found that male patients with BD and SZ harboring low-activity 5-HTTLPR alleles showed higher CpG3 DNA methylation than those harboring allele with high promoter activity. Furthermore, *in vivo* brain imaging analysis revealed a negative correlation between CpG3 DNA methylation and left amygdala volume in patients harboring low-activity alleles, suggesting a pathophysiological role of *SLC6A4* hypermethylation.

## Materials and methods

### Samples

All subjects were unrelated to each other and were ethnically Japanese. We used genomic DNA of PBCs derived from patients with SZ (N = 440) and age-matched controls (CTs) (N = 488). We also used genomic DNA of PBCs derived from an independent SZ group (N = 100), a first-episode SZ (FESZ) group (N = 16), and a group of the same number of age-matched CTs. The details of the samples before and after quality control, which involved removal of subjects with low signal intensity in pyrosequencing or with genotyping errors, are summarized in **supplementary table S1**. Details of selection criteria for FESZ and CTs were described in **supplementary methods**.

### Animals

Six adult male common marmosets (CLEA Japan, Inc., Tokyo, Japan) were used to test the effects of antipsychotics. The detailed methods have been previously described^33^ and are described in the **supplementary methods**. In brief, three marmosets were administered risperidone at a dose of 0.1 mg/kg (Wako Chemical, Tokyo, Japan), and the other 3 were given only vehicle. The substances were administered orally once a day for 28 days. All experiments were approved by the Institutional Animal Care and Use Committee and were conducted in accordance with the guidelines of the Central Institute for Experimental Animals (CIEA, Kanagawa, Japan), which comply with the Guidelines for Proper Conduct of Animal Experiments published by the Science Council of Japan Animal Care.

### Molecular methods

Detailed descriptions of the following molecular methods can be found in the **supplementary methods**: DNA preparation from PBCs of humans and animals, bisulfite modification of genomic DNA, pyrosequencing assays, 5-HTTLPR genotyping, and luciferase reporter assays for the 5-HTTLPR allele and for an *in vitro-*methylated CpG construct.

### Brain imaging analysis

Brain images were acquired using 1.5- and 3.0-tesla MRI scanners (Signa Horizon 1.5 T, Signa HDxt 3.0 T; GE Healthcare, Milwaukee, WI, USA) at the University of Tokyo Hospital as part of the Cognitive Genetics Collaborative Research Organization (COCORO) consortium.^34^ In this study, 41 CTs (32 males and 9 females) and 57 SZ patients (33 males and 24 females; 3 first-episode and 54 chronic) who had valid DNA methylation and 5-HTTLPR genotyping data were chosen for further analysis. A detailed description of the method can be found in the **supplementary methods**.

### Statistics

Comparisons of DNA methylation levels were conducted using a nonparametric Mann-Whitney U-test. The DNA methylation level in marmoset was independently measured 5 times, and the trimmed means were compared using Welch’s t-test. Differences in luciferase activity among the constructs were evaluated using the Tukey-Kramer test. The correlation between the DNA methylation and the amygdala volume was assessed with Pearson’s correlation. All statistical comparisons were two-sided. Statistical significance was set at *P* < 0.05.

## Results

### Hypermethylation of the CpG island shore of *SLC6A4* in male patients with BD

We previously identified hypermethylation at two CpG sites (named CpG3 and CpG4) within the CpG island shore of *SLC6A4* in BD using LCLs^31^ (figure 1A). Because the sample sizes were relatively small and because LCLs contain artificial epigenetic modifications,^35,36^ we tried to replicate our previous findings in a large number of PBCs from patients with BD (N = 447) and from CTs (N = 454) by pyrosequencing. We validated hypermethylation at one of the two CpG sites, CpG3 (*P* = 3.84×10^-4^, **supplementary table S2**). Consistent with a previous report,^27^ we observed apparent DNA methylation differences between male and female subjects (**supplementary figure S1**). Subgroup analysis considering sex revealed significant hypermethylation at CpG3 in male patients with BD (*P* = 0.003) but not in female patients with BD (figure 1B, **supplementary table S2**).

**Fig. 1.**
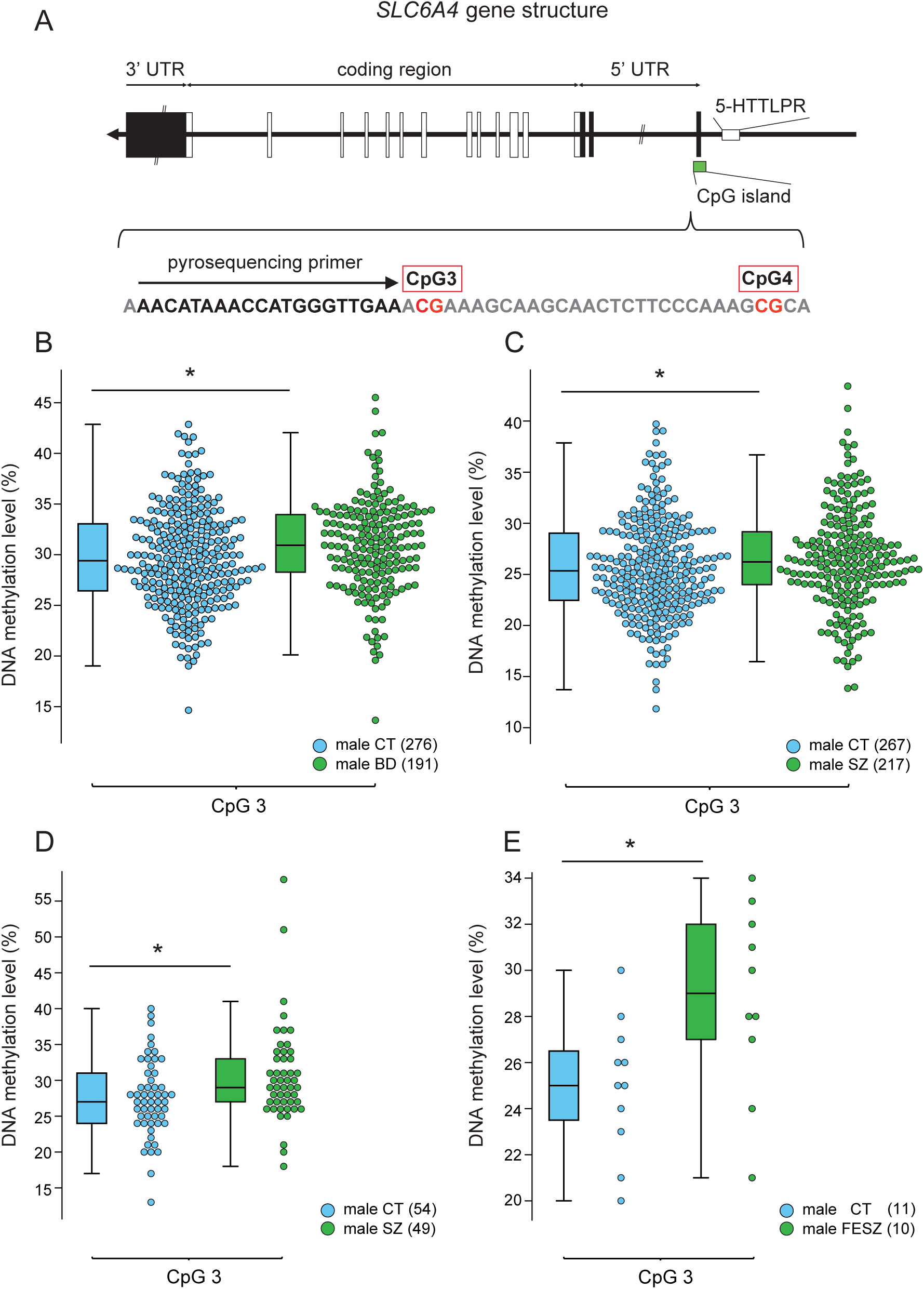
Altered DNA methylation of *SLC6A4* in male patients with BD and SZ. (**A**) Gene structure of *SLC6A4* and the targeted CpGs (CpG3: chr17:30,235,246-30,235,247 and CpG4: chr17:30,235,271-30,235,272). Comparison of DNA methylation levels was performed between male CTs and male patients with BD (**B**), SZ (set 1) (**C**), SZ (set 2) (**D**), or FESZ (**E**). The number of subjects is given in parentheses. \**P* < 0.05. 5-HTTLPR: serotonin transporter-linked polymorphic region, CT: control, BD: bipolar disorder, SZ: schizophrenia, FESZ: first-episode schizophrenia.

### Hypermethylation in PBCs of male patients with SZ

We then examined the DNA methylation levels at two CpG sites of *SLC6A4* in PBCs from patients with SZ (N = 407) and from CTs (N = 468) (set 1). Similar to the case for BD, hypermethylation at CpG3 was also found in male patients with SZ (figure 1C, **supplementary table S3**). Male-specific CpG3 hypermethylation was robustly replicated in the independent group of patients with SZ (set 2, figure 1D, **supplementary table S4**) and in the group of patients with FESZ (figure 1E, **supplementary table S5**).

### Effects of antipsychotics on DNA methylation at CpG3 in common marmosets

To assess the effects of antipsychotics, we examined CpG3 methylation levels in the blood of common marmosets treated chronically with risperidone. Notably, the genomic context around CpG3, but not CpG4, was found to be evolutionarily conserved among primates but not among rodents (**supplementary figure S2**). CpG3 DNA methylation levels were not detectable in the risperidone-treated group (N = 3) compared to the control group (N = 3, mean ± SD: 4.4 ± 0.7%).

### *In vitro* DNA methylation at CpG3 represses promoter activity

We next performed a luciferase reporter assay using constructs containing the sequences around the CpG3 region in the rat serotonergic RN46A cell line.^37^ We detected significant promoter activity around the CpG3 region. This promoter activity was abolished upon introduction of CpG3 DNA methylation (*P* < 0.01, Tukey-Kramer test) (figure 2).

**Fig. 2.**
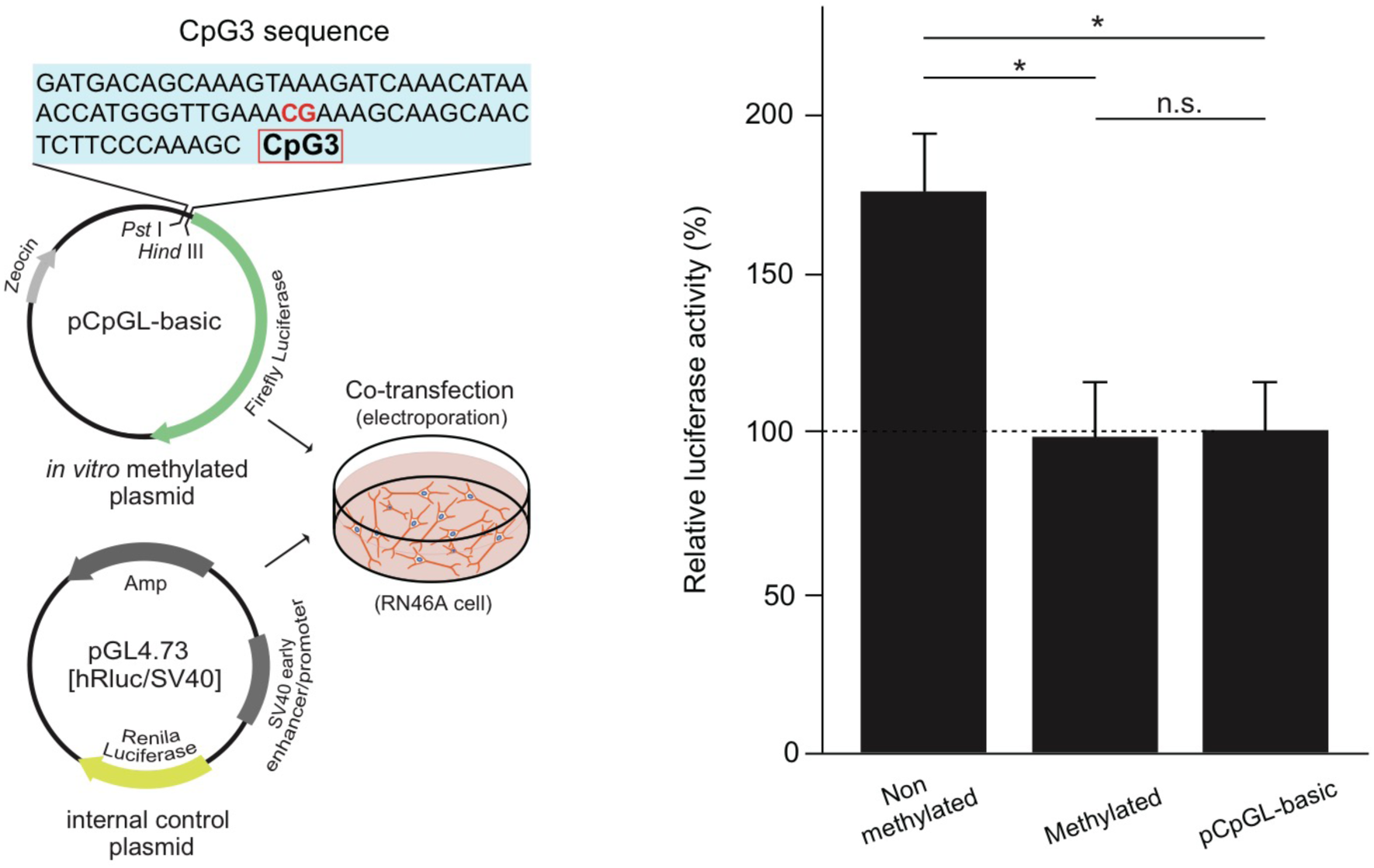
Luciferase promoter assay of a construct containing *in vitro-*methylated CpG3. *Left, in vitro*-methylated (or unmethylated) plasmid constructs including the CpG3 sequence were cotransfected with internal control plasmids into RN64A cells. *Right*, DNA methylation at CpG3 suppressed promoter activity. \**P* < 0.01. n.s.: not significant.

### Genotyping of 5-HTTLPR and promoter assay of the Asian-specific L allele

To assess the effect of 5-HTTLPR on DNA methylation, we genotyped 5-HTTLPR in CTs and patients with BD and SZ (set 1) in detail. Consistent with a previous report on a Japanese population,^2^ the subjects showed a complex distribution of allele frequencies (AFs) (**supplementary table S6**).^2^ In addition to the predominant S allele (AF: 75.0%), there were three evenly distributed L alleles: L_A_, L_G_, and L_16-C_ (AFs: 8.1%, 6.9%, and 6.4%, respectively). It should be noted that L_16-C_ has not been reported in the Caucasian population. Given that L_A_ has the highest promoter activity and that L_G_ shows low promoter activity equal to that of S_A_,^^38,39^^ we determined the promoter activity of L_16-C._ We found that L_16-C_ showed low promoter activity, similar to S_A_ (**supplementary figure S3**).

### Patients harboring low-activity 5-HTTLPR alleles showed high DNA methylation levels at CpG3

We then conducted a case-control DNA methylation analysis considering the 5-HTTLPR genotype. Consistent with previous reports,^24,31^ we observed significant hypermethylation in patients with SZ harboring homozygous S_A_ compared to CTs with the same alleles (*P* = 0.040, table 1). While patients harboring L_A_ did not show any DNA methylation differences, those harboring L_16-C_ or L_G_ showed hypermethylation compared to CTs (*P* = 0.049). Given that only L_A_ showed high promoter activity, while the others showed similar levels of low activity, we then conducted a promoter activity-based case-control analysis. We found that patients with low-activity alleles (S_A_, L_16-C_, or L_G_ alleles) showed more robust hypermethylation than CTs (*P* = 0.006, table 1). The same relationship was also maintained in BD, though the effect of the low-activity alleles seemed to be different from that in SZ, likely due to the limited sample sizes (**supplementary table S7**). We examined CpG3 DNA methylation in females in the same datasets but did not find any significant differences in patients with SZ or BD compared to CTs (**supplementary table S8**).

**Table 1.** Case-control DNA methylation analysis of SZ considering the promoter activity of 5-HTTLPR.

| Allele<br>(S<sub>A</sub> background) | Diagnosis | N | CpG 3 |  |
| --- | --- | --- | --- | --- |
|  |  |  | DNA methylation<br>level (%)<br>(mean±SD) | *P*-value<br>(Cohen's d) |
| S<sub>A</sub> | CT | 149 | 25.5±4.9 | **0.040** |
|  | SZ | 145 | 26.5±5.0 | (0.200) |
| L<sub>A</sub> | CT | 34 | 26.2±4.2 | 0.822 |
|  | SZ | 21 | 26.3±4.5 | (0.031) |
| L<sub>16-C</sub>/L<sub>G</sub> | CT | 50 | 25.1±4.3 | **0.049** |
|  | SZ | 37 | 26.8±5.2 | (0.377) |
| S<sub>A</sub>/L<sub>16-C</sub>/L<sub>G</sub><br>(All low-activity<br>alleles) | CT | 199 | 25.4±4.7 | **0.006** |
|  | SZ | 182 | 26.6±5.0 | (0.240) |

CT: control, SZ: schizophrenia, 5-HTTLPR: serotonin transporter-linked polymorphic region. Significant *P*-values are shown in bold.

### Data from patients with SZ harboring low-activity 5-HTTLPR alleles showed a negative correlation between DNA methylation levels and amygdala volumes

The amygdala is essential for emotional processing, and its activation is thought to be associated with 5-HTTLPR.^40^ Recently, large-scale cross-sectional studies have revealed that amygdala volumes are significantly reduced in patients with SZ compared to healthy subjects.^34,41^ Furthermore, previous studies have reported an association between *SLC6A4* promoter region methylation and increased threat-related activity in the left amygdala.^20,42^ Using three-dimensional brain images from our samples, we found that CpG3 DNA methylation levels were negatively correlated with left amygdala volumes in male patients with SZ harboring low-activity alleles (n = 24, Pearson’s *R* = -0.454, *P* = 0.026) but not in male CTs (n = 26, Pearson’s *R* = -0.099, *P* = 0.631) (figure 3). A significant negative correlation was observed only in male patients with SZ (**supplementary figure 4**).

**Fig. 3.**
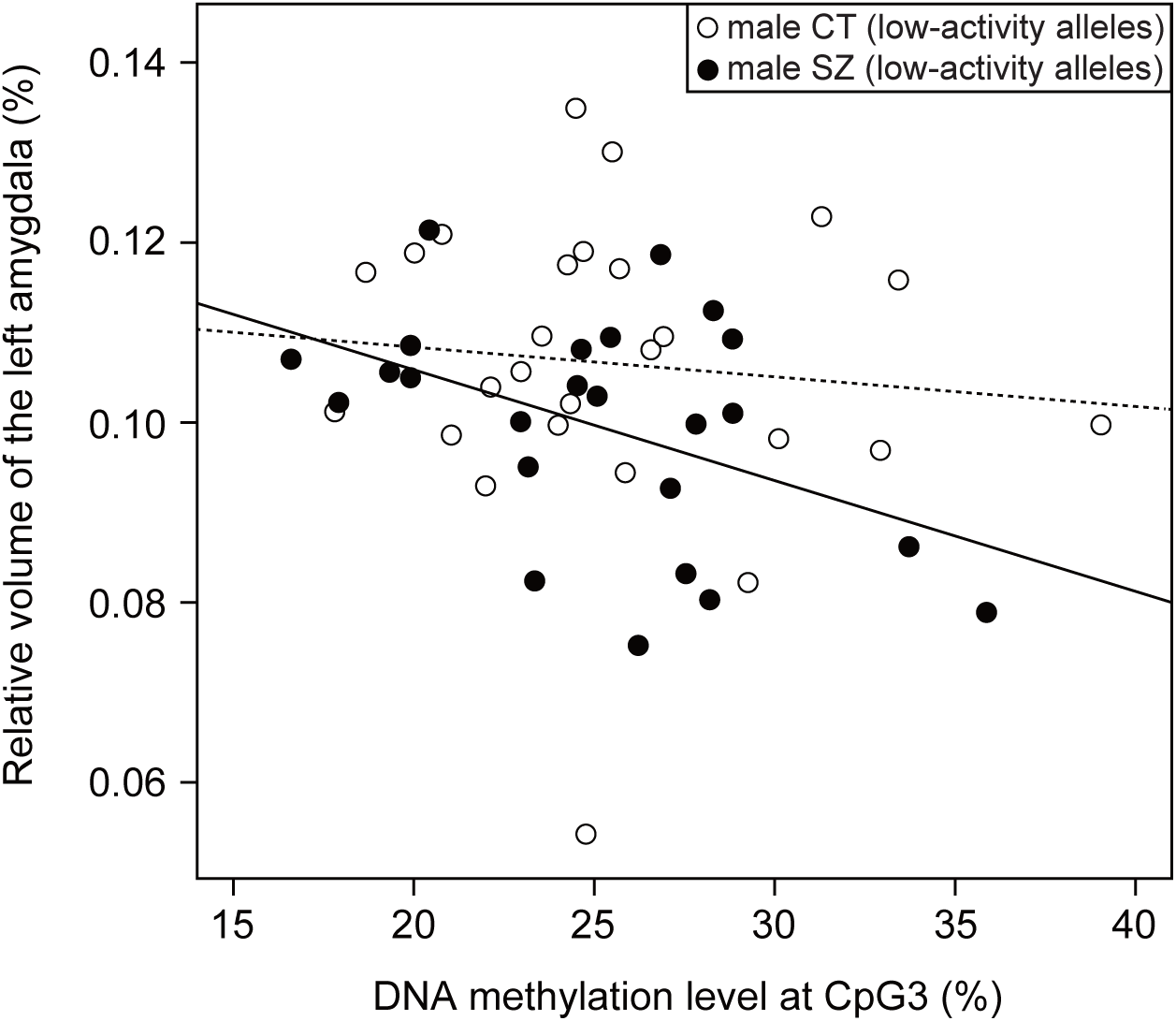
Correlation between DNA methylation levels at CpG3 and left amygdala volumes. DNA methylation levels at CpG3 were negatively correlated with left amygdala volumes in male patients with SZ carrying low-activity alleles (solid line) but not in male CTs (broken line). CT: control, SZ: schizophrenia.

## Discussion

In this study, we identified hypermethylation of the promoter region of *SLC6A4* in blood samples from male patients with SZ. Hypermethylation was evident in male patients with low-activity 5-HTTLPR alleles and was inversely correlated with amygdala volume. It should be noted that we could not apply multiple testing corrections in our analyses because performing stratified comparisons reduced the sample sizes. Instead, we used several independent sample groups to corroborate the results.

### Effects of sex differences, age and medication

Based on the apparent DNA methylation differences between males and females (**supplementary figure S1**), we decided to separately analyze DNA methylation levels with regard to sex rather than including sex as a covariate. Previous studies have also reported higher DNA methylation in females than males at the CpG island or CpG island shore of *SLC6A4* in cord blood,^43-45^ PBCs,^46^ and postmortem prefrontal cortices.^47^ Although the underlying mechanism of the elevated DNA methylation in females remains elusive, choline, a precursor for the methyl group donor betaine, might be a key molecule given the enhanced endogenous reserves and *de novo* biosynthesis of choline by estrogen in females.^48,49^ Indeed, dietary choline supplementation or deficiency in rats results in global and site-specific DNA methylation changes in metabolic-related and neural development-related genes,^50-52^ and some of these differences are sex-dependent.^51^ The reason for the allele-dependent DNA methylation differences between patients of different sexes also remains unclear. The significant hypermethylation of *SLC6A4* in male patients suggests that this modification may be associated with differences in the clinical features and courses of psychosis between males and females.^53^

To analyze the contributions of age and diagnosis to CpG3 DNA methylation status, we conducted a multiple linear regression analysis and identified significant contributions or tendencies toward contributions of diagnosis and age in the male BD-control analysis (*P* = 0.011), the male SZ-control (set 1) analysis (*P* = 0.011), and the male SZ-control (set 2) analysis (*P* = 0.054). However, no significant contributions were detected in females. These observations indicate that CpG3 DNA methylation status is modulated not only by diagnosis but also by age in males.

Age of disease onset may be a confounding factor for DNA methylation changes. To address this issue, we examined the DNA methylation levels of male SZ patients separated by age of disease onset: one group included patients less than 20 years old (the “under-20 group”; n = 57, average onset age = 16.9 ± 1.5), and the other included patients from 20 to less than 40 years old (the “over-20 group”; n = 134, average onset age = 27.1 ± 4.7). The average age at blood sampling was matched between the two groups (under-20 group = 43.8 ± 11.7, over-20 group = 44.0 ± 10.4). The results revealed that the DNA methylation levels at CpG3 did not significantly differ between the two groups (under-20 group = 26.8 ± 5.4, over-20 group = 26.6 ± 4.8), suggesting that the age of onset did not affect the DNA methylation changes at CpG3.

Regarding medication, we tested the influence of antipsychotics on the DNA methylation level of CpG3 using PBCs from common marmosets and found decreased rather than increased DNA methylation in the PBCs of antipsychotic-treated marmosets compared to those of control animals. A similar decrease in DNA methylation of CpG3 was previously reported in a cell line cultured with mood stabilizers.^54^ These findings suggest that medication is unrelated to the increased DNA methylation at CpG3 in patients. These results also suggest that medication may play a role in decreasing CpG3 methylation levels, though further mechanistic studies are needed to address this possibility. It should be noted that the number of animal samples was too small to reach a conclusion and that these findings should be independently replicated.

### Complexity of 5-HTTLPR and its association with DNA methylation

In this study, S_A_ was present at a significantly higher frequency in cases than in controls (SZ-BD vs CT: *P* < 0.001, SZ vs CT: *P* < 0.008, BD vs CT: *P* < 0.003, Fisher’s exact test, **supplementary table S6**). This finding is consistent with previous reports.^5-8^ However, careful interpretation is needed because a recent large-scale studies did not support such associations with 5-HTTLPR.^12,13^

Previous studies have revealed that DNA methylation changes in *SLC6A4* are associated with childhood abuse, stressful life events, and depression in S allele carriers.^15,21,24,26,55^ Consistent with these findings, rhesus macaques harboring the S allele exhibit higher DNA methylation levels in *SLC6A4* and lower *SLC6A4* expression in PBCs when exposed to maternal or social separation than those not harboring the S allele.^56^ However, other studies have failed to identify S allele-linked DNA methylation changes in depressed subjects.^22,25,30^ These inconsistent findings could partly stem from the complexity of 5-HTTLPR genotypes, which results in variable *SLC6A4* promoter activity. Hu *et al*^38^ revealed that the L_G_ allele, which is the minor L allele with a G substitution (rs25531), exhibits lower promoter activity than the major L allele with an A substitution (L_A_) and that the S and L_G_ alleles exhibit nearly equivalent promoter activity. In addition to L_G_, we revealed that L_16-C_, one of the major L alleles in the Asian population^^2,57^^ but not in the European population (**supplementary table S6**), has low promoter activity similar to that of the S and L_G_ alleles (**supplementary figure S3**).

We found that low-activity 5-HTTLPR alleles were robustly associated with higher DNA methylation levels in patients through a promoter activity-based case-control association approach. Our findings will be important for future 5-HTTLPR studies across different ethnic populations and may contribute to understanding the contradictory results of 5-HTTLPR genetic association studies. The molecular mechanism that connects genotype to DNA methylation status remains unclear. One hypothesis involves the participation of the G-quadruplex within 5-HTTLPR,^58^ which is a four-stranded noncanonical B-form DNA structure and is known to interact directly with DNA nucleotide methyltransferase enzymes.^59^

### Functional and pathophysiological role of CpG3 DNA methylation

We confirmed the existence of CpG3 hypermethylation in a group of male patients with SZ (set 1), an independent group of patients with SZ (set 2), a group of patients with FESZ and a group of patients with BD (**supplementary tables 2, 3, 4, and 5**), implicating the existence of common DNA methylation changes in individuals with psychosis. Hypermethylation was found in both the first episode and in chronic stages of SZ, suggesting that this epigenetic change occurs at an early stage and lasts for a long time during the course of the illness. Future studies will include longitudinal assessments of patients with consideration of their ratings of symptom severity.

Associations between *SLC6A4* DNA methylation and *SLC6A4* expression have previously been reported,^16,27,42,60,61^ and we previously demonstrated an inverse correlation between CpG3 DNA methylation levels and *SLC6A4* expression levels in LCLs.^31^ In this study, we could not assess *SLC6A4* expression levels in PBCs because of the unavailability of RNA samples. However, we found that DNA methylation of CpG3 was sufficient for gene silencing in a cell culture model using a CpG3-specific *in vitro*-methylated reporter.

Although the DNA methylation changes at CpG3 in patients were subtle, our *in vitro* luciferase assays suggested the importance of the patterns of methylated alleles. If two CpG3 alleles in a cell are methylated, that cell does not express *SLC6A4* at all. Elevations in the numbers of such cells may cause more dysfunction than elevations in the numbers of cells with only one methylated allele, even though the apparent DNA methylation changes seemed to be subtle.

The genomic region around CpG3 is conserved among primates but not among rodents (**supplementary figure S2**). Using the JASPAR transcription factor binding profile database (http://jaspar.genereg.net/) and the Genotype-Tissue Expression (GTEx) gene expression database (https://www.gtexportal.org/), we found that the putative binding sites of Elk-1 (*ELK1*) and Meis homeobox 3 (*MEIS3*), which are highly expressed in the brain, were predicted to overlap with the sequence containing CpG3 (data not shown). Interestingly, *MEIS3* is strongly expressed in the amygdala and anterior cingulate cortex, suggesting a possible link between the function of the CpG3 site and the development of the primate brain.

Recent large-scale cross-sectional analyses have revealed significant alterations in amygdala volume in patients with SZ.^34,41^ Previous studies have suggested that S allele carriers have smaller amygdala volumes than L homozygotes do.^62-64^ Another study reported reduced volumes and excessive threat-related reactivity in the amygdalae of depressed patients.^65^ In addition, two studies have shown increased threat-related left amygdala reactivity is positively correlated with DNA methylation changes in *SLC6A4*.^20,42^ Taken together, these findings suggest that amygdala volume may be associated with DNA methylation in *SLC6A4* and 5-HTTLPR alleles. We therefore specifically examined this association and found a significant inverse correlation between CpG3 DNA methylation levels and left amygdala volumes in male patients with SZ harboring low-activity alleles (figure 3). Although it remains unclear about the mechanisms by which DNA methylation status in PBCs correlates with the brain volume, a recent epigenome-wide meta-analyses indicated their relationship.^66^ Besides, previous findings that LCLs and postmortem brain from patients with BD share a common CpG3 hypermethylation^31^ and that *SLC6A4* DNA methylation status in amygdala tissue is indeed associated with its expression^42^ could support our observations. Given that increases in DNA methylation of *SLC6A4* are followed by increases in threat-related left amygdala reactivity,^20^ the reduced amygdala volumes we observed may have been accompanied by altered threat-related reactivity in the patients. More comprehensive studies involving large-scale *in vivo* imaging experiments are needed to elucidate the effects of altered DNA methylation on other brain regions.

### Considerations of cell type and blood-brain correlations

Blood cell type affects DNA methylation status^67^ and has been proposed to be a confounding factor for *SLC6A4* DNA methylation.^68^ In this study, we could not consider the effect of blood cell composition. However, considering that CpG3 hypermethylation was initially identified and replicated in LCLs,^31^ which are established from B cells, blood cell type may not be a major confounding factor in this study. Besides, we confirmed that CpG3 hypermethylation affected *SLC6A4* promoter activity in the rat serotonergic cell line, implicating the blood-brain DNA methylation correlations for CpG3. However, cell type-specific analysis is needed to determine whether other cell types have common DNA methylation changes.

## Conclusion

We report that male patients with SZ and BD harboring low-activity 5-HTTLPR alleles exhibited increased DNA methylation levels at the CpG island shore of *SLC6A4*. This epigenetic change started at a very early psychotic stage and could associate with reduced amygdala volume via *SLC6A4* downregulation. Further mechanistic studies and *in vivo* imaging studies would be useful for elucidating the pathophysiology of epigenetic alteration of *SLC6A4*.

## Supplementary Material

Supplementary data are available at *Schizophrenia Bulletin* online.

## Acknowledgements

We thank H. Saida and T. Miyauchi for their technical assistance. We also thank M. Suga, H. Inoue, Y. Aoki, H. Takao, H. Sasaki, and H. Yamada for their recruitment of subjects, brain imaging data acquisition, and data management. Some of the computations were performed at the Research Center for Computational Science, Okazaki, Japan.

