## Supplemental data for "Promoter activity-based case-control association study on *SLC6A4* highlighting hypermethylation and altered amygdala volume in male patients with schizophrenia"

### Supplementary Methods

#### Samples

We used genomic DNA of peripheral blood cells (PBCs) derived from patients with bipolar disorder (BD) (N = 450) as well as age-matched controls (CT) (N = 488). Demographics were summarized in **supplementary table S1**. The subjects were unrelated each other and ethnically Japanese. Every subject received a detailed description of this study and provided written informed consent. All patients were diagnosed according to the DSM-IV (Diagnostic and Statistical Manual of Mental Disorders, Fourth Edition) criteria by at least two experienced psychiatrists. The ethics committees of the Kumamoto University and collaborative research organizations approved this study.

FESZ patients were identified according to the following criteria: (i) they had experienced a first acute psychosis as defined by the Structured Interview for Prodromal Symptoms (SIPS) and were diagnosed with SZ according to the DSM-IV criteria after 6 months of follow-up, (ii) they had received antipsychotic medication for less than 16 cumulative weeks, and (iii) their continuous psychotic symptoms had lasted for 60 months or less. CTs were selected from among voluntarily recruited employees, students, and their friends at the University of Tokyo Hospital and were subjected to unstructured interviews by senior psychiatrists (N = 91), the Mini-International Neuropsychiatric Interview (MINI) (N = 430), and the Composite International Diagnostic Interview (CIDI) (N = 9). All participants were confirmed to have (i) no current or past Axis I psychiatric or physical diagnoses and (ii) no first-degree relatives with SZ.

**Molecular methods**

**Animals**

Six adult male common marmosets (CLEA Japan, Inc., Tokyo, Japan) were used for testing the effect of antipsychotics. Three of them (medicated marmosets: mean age ± SD = 3.78 ± 0.47) were administered risperidone at 0.1 mg/kg (Wako Chemical, Tokyo, Japan) suspended in 2 mL/kg of 0.5% methylcellulose as vehicle (Wako Chemical), and the other 3 (CT marmosets: mean age ± SD = 3.38 ± 0.46) were given only vehicle. These were administered orally once a day. After 28 days of drug administration, all subjects were sacrificed by injecting ketamine (10-30 mg/kg) and xylazine (0.6-2.4 mg/kg), followed by isoflurane (3-5%) inhalation. PBCs were collected in EDTA-coated tubes and centrifuged for 10 min at 11,500 *g*. The centrifuged PBC components were hemolyzed with 0.2% NaCl and stored at -80°C.

**DNA extraction and bisulfite modification**

DNA extraction was performed using a Wizard Genomic DNA Purification Kit (Promega, Madison, WI, USA) on whole blood and blood cells or using a Genomix DNA isolation kit (Talent, Trieste, Italy) on the buffy coat following the manufacturer's instructions. Common marmoset genomic DNA from whole blood was purified using phenol/chloroform extraction. Two micrograms of human genomic DNA and 500 ng of common marmoset genomic DNA were treated with sodium bisulfite conversion using an EpiTect 96 Bisulfite Kit (QIAGEN, Valencia, CA, USA) following the manufacturer's protocols.

**Bisulfite PCR amplification**

The CpG island shore of *SLC6A4* (chr17:30,235,139-30,235,342 (GRCh38/hg38)) was amplified with forward primer 5'-TTTTTAGTTGTTTGGGTATTTGTGTTA-3' and 5'-biotinylated reverse primer Bio-5'-AAAACTTTACAACCTCTTAAAAACCC-3' using bisulfite-converted genomic DNA as a template. The bisulfite PCR amplification was performed in a total volume of 50 μL containing 5 μL of 10 x PCR Amplification Buffer (Invitrogen, Carlsbad, CA, USA), 10 μL of 5 M betaine (Sigma Aldrich, St Louis, MO,

USA), 3 µL of 50 mM $MgCl_2$ (Invitrogen), 1 µL of 10 mM dNTP (Invitrogen), 2 µL each of 10 µM primers, 2 ng of Single-Stranded DNA Binding Protein (Promega), 5 U of Platinum *Taq* DNA Polymerase (Invitrogen), and 1 µL of bisulfite-modified genomic DNA. The thermocycling conditions were an initial incubation at 95°C for 3 min followed by 40 cycles each of 10 s at 98°C, 30 s at 55°C, and 15 s at 72°C. For common marmosets, the target *SLC6A4* region was amplified with a forward primer (5'-TTTTTAGTTATTTGGGTATTTGTGTTA-3') and a 5'-biotinylated reverse primer (Bio-5'-AAAACTTTACAACCTCTTAAAAACCC-3'). The bisulfite PCR amplification was performed in a total volume of 25 µL containing 2 µl of bisulfite-modified common marmoset genomic DNA. The proportions of reaction components were the same as above. All PCR fragments were checked on a 2.5% agarose gel to ensure successful single-banded amplification before proceeding to pyrosequencing.

**Pyrosequencing**

Bisulfite PCR fragments of BD, SZ (set 2), FESZ, and their age-matched controls were processed for pyrosequencing analyses according to the standard protocol described in a previous study.[31] A PyroMark Q96 Vacuum Workstation (QIAGEN) was used when bisulfite

PCR fragments of SZ (set 1), common marmoset, and their age-matched controls were processed for pyrosequencing analyses (QIAGEN). The former method used a filter plate, a MultiScreen-HV, Clear Plate (Millipore, Billerica, MA, USA), while the latter used filter probes to capture PCR fragments. The method using filter probes has an advantage in shortening the processing time and purifying PCR fragments homogeneously because of simultaneous vacuum reaction. Briefly, 38 μL of bisulfite PCR products (22 μL of common marmoset bisulfite PCR products and 16 μL of water) were mixed with 2 μL of streptavidin-sepharose beads (Amersham Biosciences) and 40 μL of PyroMark Binding Buffer (QIAGEN) on a 96-well PCR plate, and the plate was vortexed for 10 min at room temperature. Next, the vacuum tool's filter probes were inserted into the plate, and all beads were captured. The filter probes were transferred through 100 mL of 70% ethanol for 5 s, denaturation solution (0.2 N NaOH) for 5 s, and washing buffer for 10 s. Finally, the captured beads were released into a PSQ 96 Plate (QIAGEN) containing 20 μL of PyroMark Annealing Buffer (QIAGEN) with 1.5 μL of 10 μM sequencing primer (5'-AATATAAATTATGGGTTGAA-3' for human or 5'-AATATAAATTAAGGGTTGAA-3' for common marmoset) by gently shaking the tool in the wells. This plate was then heated to 96°C for 2 min. Processed PCR fragments were used for pyrosequencing on a PSQ 96MA instrument (QIAGEN) according to the manufacturer's instructions. The positions of the measured CpG sites were chr17:30,235,246-30,235,247 for CpG3 and chr17:30,235,271-30,235,272 for CpG4 (GRCh38/hg38). DNA methylation levels of CpG3 and CpG4 were measured in a single pyrosequencing reaction with one sequencing primer. We confirmed the accuracy and linearity of pyrosequencing over a wide range of DNA methylation levels at CpGs using bisulfite PCR products that were amplified from mixtures of *in vitro* methylated and unmethylated DNA. The DNA methylation level of each CpG was calculated with PSQ 96MA software (QIAGEN). Statistical analyses were performed in the R software package (R Foundation for Statistical Computing, Vienna, Austria).

DNA methylation values of each sample set were measured independently in a different period of time. Due to the slight differences of pyrosequencing procedure in each sample set, we did not compare the DNA methylation values across different sample sets.

**5-HTTLPR genotyping**

5-HTTLPR was amplified with 5'-GGTGAAATTCCCAAGCTTGTTG-3' and 5'-TTCTGGTGCCACCTAGACGC-3' as the primers. PCR amplification was performed in a total volume of 20 μL containing 4 μL of 5 x Promega Flexi buffer (Promega), 4.8 μL of 5 M betaine (Sigma Aldrich), 1.6 µL of 25 mM $MgCl_2$ (Promega), 0.8 µL of 10 mM dNTP mix (Invitrogen), 4 µL each of 10 µM primers, 0.75 U of GoTaq Hot Start DNA Polymerase (Promega), and 50 ng of genomic DNA. The thermocycling conditions were an initial incubation at 94°C for 2 min; 33 cycles each of 30 s at 94°C, 30 s at 65°C, and 30 s at 72°C; and 1 cycle of 10 min at 72°C. The alleles were confirmed by electrophoresis in 2.5% agarose gel, and DNA sequencing was performed in both directions using 5'-TTGTTGGGGATTCTCCCG-3' or 5'-CTGGTGCCACCTAGACGCC-3' as a sequencing primer.

**Plasmid construction**

The CGI shore containing CpG3 but not CpG4 or other CpG sites (chr17:30,235,203-30,235,271 (GRCh38/hg38)), named the CpG3 sequence, was amplified with 5'-AActgcagCTTTGGGAAGAGTTGCTTGCTT-3' with a *Pst*I site and 5'-AAaagcttGATGACAGCAAAGTAAAGATC-3' with a *Hind*III site as the primers. PCR amplification was performed in a total volume of 25 µL containing 2.5 µL of 10 x PCR buffer for KOD-Plus-Neo (Toyobo, Osaka, Japan), 1.5 µL of 25 mM $MgSO_4$ (Promega), 0.5 µL of 10 mM dNTP mix (Invitrogen), 0.75 µL each of 10 µM primers, 0.5 U of KOD-Plus-

Neo (Toyobo), and 50 ng of genomic DNA. The thermocycling conditions were an initial incubation at 94°C for 2 min and 33 cycles each of 10 s at 98°C and 15 s at 65°C. The PCR fragments were gel-purified with a MinElute Gel Extraction Kit (Promega) followed by 3' A-attachment with TaKaRa ExTaq (Takara, Tokyo, Japan) and cloned into the pCR2.1 vector using the TOPO cloning kit (Invitrogen) with DH5α Competent Cells (Takara). A single bacterial colony was cultured and purified with a Wizard Plus Minipreps DNA Purification System (Promega). The inserted DNA fragment was extracted with *Pst*I and *Hind*III (New England Biolabs, MA, USA) and then subcloned into a CpG-free pCpGL-basic firefly luciferase reporter vector, kindly provided by Prof. Dr. Michael Rehli (University Hospital Regensburg, Germany), with DH5α cells. A single bacterial colony was cultured and purified with an endotoxin-free plasmid DNA purification kit (NucleoBond Xtra EF purification system, Macherey-Nagel GmbH, Düren, Germany). Plasmid constructs were sequenced to verify the proper ligation and absence of artificial mutations. To create *in vitro* methylated constructs, 20 μg of the construct were treated with *Sss*I DNA methyltransferase (New England Biolabs) according to the manufacturer's protocol and purified with a Wizard SV Gel and PCR Clean-Up System (Promega). The unmethylated construct was prepared similarly to methylated construct, but without adding *Sss*I. The 5-HTTLPR variants ($S_A$ and

$L_{16\text{-}C}$) were amplified with 5’-AAgagctcGGTGAAATTCCCAAGCTTGTTG-3’ with a *Sac*I site and 5’-AActcgagTTCTGGTGCCACCTAGACGC-3’ with a *Xho*I site as primers. The reaction volume and compositions of PCR were the same as described in the plasmid construction section. The thermocycling conditions were an initial incubation at 94°C for 2 min and 32 cycles each of 10 s at 98°C and 30 s at 72°C. The PCR fragments were cloned into the pCR2.1 vector and then subcloned into the pGL4.10 firefly luciferase reporter vectors (Promega) using the *Sac*I and *Xho*I sites (New England Biolabs). A single bacterial colony was cultured and purified as described above. All constructs were sequenced to verify the proper ligation and absence of artificial mutations.

**Luciferase reporter assay**

Undifferentiated rat raphe-derived cell lines (Sigma Aldrich) were grown in D-MEM/Ham's F-12 medium (Wako Chemical) supplemented with 10% fetal bovine serum (SAFC Bioscience, Lenexa, KS, USA) and 0.25 mg/mL of G418 (Geneticin, Life Technologies, Gaithersburg, MD, USA) at 5% $CO_2$ and their permissive temperature of 33°C. The cells were transfected with 2 ug of either luciferase reporter construct or a positive control vector, pCMV-EGFP (NEPA GENE, Tokyo, Japan), and 200 ng of pGL4.73 *Renilla* luciferase reporter vector (Promega) as an internal control using a NEPA21 electroporator (NEPA GENE). Twenty-four hours after transfection, the cells were harvested and lysed. Luciferase activity was measured using a Dual Luciferase Assay System (Promega) and a GloMax™ 96 Microplate Luminometer (Promega) according to the manufacturer's protocol. Each construct was transfected in triplicate, and each assay was independently performed in triplicate. Firefly/Renilla ratios were calculated in each transfection and normalized to the mean ratio of blank control luciferase vector (pGL4.10 or pCpGL-basic, Promega).

***In vivo* imaging analysis**

For volumetric analysis, the 1.5-tesla MRI used a spoiled gradient echo (SPGR) sequence and a circularly polarized head coil (repetition time = 35 ms, echo time = 7 ms, flip angle = 30°, matrix = 256 x 256 x 124, field of view = 186 x 240 x 240 mm, voxel size = 0.9375 x 0.9375 x 1.5 mm, slice thickness = 1.5 mm, number of axial slices = 124). The 3.0-tesla MRI used a fast SPGR sequence and an 8-channel coil (repetition time = 6.8 ms, echo time = 1.94 ms, flip angle = 20°, matrix = 256 x 256 x 176, field of view = 176 x 256 x 256 mm, voxel size = 1 x 1 x 1 mm, slice thickness = 1 mm, number of axial slices = 176). After removing low-quality MRI images (e.g., those with low signal-to-noise ratios, motion artifacts, magnetic susceptibility artifacts, or abnormal organic findings), the remaining images were processed with FreeSurfer software version 5.3 (http://surfer.nmr.mgh.harvard.edu) to obtain the brain's subcortical volumes and intracranial volume (ICV). The ratio of left amygdala volume to intracranial volume was adjusted using a linear regression analysis including sex, age, and MRI scanner types as covariates. The linear regression analysis was performed in R.

**Supplementary Table S1.** Demographics of analyzed sample sets

| Diagnosis | N | Age | Sex (male:female) | Male age | Female age |
|---|---|---|---|---|---|
| CT | 454 (460) | 45.3±10.7 (45.2±10.7) | 276:178 (280:180) | 45.3±10.2 (45.2±10.2) | 45.2±11.5 (45.2±11.5) |
| BD | 447 (450) | 46.2±13.1 (46.2±13.1) | 191:256 (194:256) | 46.1±10.7 (46.1±10.8) | 46.3±14.6 (46.3±14.6) |
| CT | 468 (488) | 45.2±10.6 (45.2±10.5) | 267:201 (280:208) | 45.1±10.3 (45.2±10.2) | 45.3±11.0 (45.2±10.9) |
| SZ (set 1) | 407 (440) | 45.2±11.2 (45.3±11.1) | 217:190 (228:212) | 45.2±11.3 (45.2±11.1) | 45.1±11.1 (45.4±11.2) |
| CT | 99 (100) | 42.2±12.0 (42.2±12.0) | 54:45 (55:45) | 42.8±12.6 (42.8±12.5) | 41.4±11.3 (41.4±11.3) |
| SZ (set 2) | 94 (100) | 43.1±13.2 (43.1±13.0) | 49:45 (54:46) | 43.8±14.0 (43.8±13.8) | 42.4±12.3 (42.3±12.2) |
| CT | 16 | 22.8±4.3 | 11:5 | 22.3±5.0 | 24.0±2.3 |
| FESZ | 16 | 21.8±4.9 | 10:6 | 21.5±5.6 | 22.2±3.7 |

CT: control, BD: bipolar disorder, SZ: schizophrenia, FESZ: first episode schizophrenia. The numbers in the parentheses indicates the initial samples before quality control. All case-control studies were performed independently in different periods of time. Note that CTs were overlapped across sample sets: The number of CTs originally collected for BD analysis was 460. For SZ analysis, 28 female CTs were added to this sample set.

**Supplementary Table S2.** DNA methylation difference of *SLC6A4* in BD

| | | | CpG 3 | | CpG 4 | |
|---|---|---|---|---|---|---|
| Sex | Diagnosis | N | DNA methylation level (%) (mean±SD) | *P*-value (Cohen's d) | DNA methylation level (%) (mean±SD) | *P*-value (Cohen's d) |
| All | CT | 454 | 31.9±5.5 | **$3.84 \times 10^{-4}$** (0.248) | 56.5±6.1 | 0.098 (0.133) |
| | BD | 447 | 33.3±5.3 | | 57.3±5.7 | |
| Male | CT | 276 | 29.7±4.9 | **0.003** (0.249) | 54.2±5.5 | 0.548 (0.071) |
| | BD | 191 | 30.9±4.6 | | 54.5±5.0 | |
| Female | CT | 178 | 35.3±4.8 | 0.350 (0.060) | 60.2±5.1 | 0.055 (0.162) |
| | BD | 256 | 35.0±5.1 | | 59.4±5.2 | |

CT: control, BD: bipolar disorder. Significant *P*-values are shown in bold.

**Supplementary Table S3.** DNA methylation difference of *SLC6A4* in SZ (set 1)

| Sex | Diagnosis | N | CpG 3 | | CpG 4 | |
|---|---|---|---|---|---|---|
| | | | DNA methylation level (%) (mean±SD) | *P*-value (Cohen's d) | DNA methylation level (%) (mean±SD) | *P*-value (Cohen's d) |
| All | CT | 468 | 28.3±5.6 | 0.114 (0.110) | 57.4±6.8 | **0.049** (0.141) |
| | SZ | 407 | 28.9±5.7 | | 58.4±7.0 | |
| Male | CT | 267 | 25.7±4.8 | **0.032** (0.176) | 54.4±6.2 | 0.197 (0.177) |
| | SZ | 217 | 26.6±5.0 | | 55.2±6.3 | |
| Female | CT | 201 | 31.7±4.7 | 0.748 (0.025) | 61.3±5.4 | 0.266 (0.118) |
| | SZ | 190 | 31.5±5.3 | | 62.0±5.9 | |

CT: control, SZ: schizophrenia. Significant *P*-values are shown in bold.

**Supplementary Table S4.** DNA methylation difference of *SLC6A4* in SZ (set 2)

| Sex | Diagnosis | N | CpG 3 | | CpG 4 | |
|---|---|---|---|---|---|---|
| | | | DNA methylation level (%) (mean±SD) | *P*-value (Cohen's d) | DNA methylation level (%) (mean±SD) | *P*-value (Cohen's d) |
| All | CT | 99 | 30.6±5.8 | 0.024 (0.397) | 57.2±6.0 | 0.082 (0.294) |
| | SZ | 94 | 33.1±6.9 | | 59.1±6.9 | |
| Male | CT | 54 | 27.5±5.4 | **0.010** (0.519) | 54.4±5.8 | 0.193 (0.290) |
| | SZ | 49 | 30.7±6.8 | | 56.1±6.3 | |
| Female | CT | 45 | 34.2±4.0 | 0.216 (0.295) | 60.6±4.3 | 0.174 (0.334) |
| | SZ | 45 | 35.7±6.0 | | 62.4±6.0 | |

CT: control, SZ: schizophrenia. Significant *P*-values are shown in bold.

**Supplementary Table S5.** DNA methylation difference of *SLC6A4* in FESZ

| Sex | Diagnosis | N | CpG 3 | | CpG 4 | |
|---|---|---|---|---|---|---|
| | | | DNA methylation level (%) (mean±SD) | *P*-value (Cohen's d) | DNA methylation level (%) (mean±SD) | *P*-value (Cohen's d) |
| All | CT | 16 | 28.0±5.5 | 0.162 (0.460) | 51.3±4.3 | 0.131 (0.464) |
| | SZ | 16 | 30.5±5.3 | | 54.1±7.4 | |
| Male | CT | 11 | 25.0±2.9 | **0.026** (1.079) | 49.5±2.9 | **0.031** (0.959) |
| | SZ | 10 | 28.8±4.1 | | 54.5±6.9 | |
| Female | CT | 5 | 34.6±3.8 | 0.926 (0.236) | 55.2±4.6 | 0.927 (0.232) |
| | SZ | 6 | 33.3±6.3 | | 53.5±8.9 | |

CT: control, SZ: schizophrenia. Significant *P*-values are shown in bold.

**Supplementary Table S6.** The allele frequencies of 5-HTTLPR in Japanese population

| Allele | AF from a previous study (Nakamura et al., Mol Psychiatry 2000) | | AF from this study | | |
|---|---|---|---|---|---|
| | Caucasian (n = 148) | Japanese (n = 262) | CT (n = 936) | SZ (n = 814) | BD (n = 894) |
| $S_{14\text{-}A}$ ($S_A$) | 66 (44.6%) | 207 (79.0%) | 702 (75.00%) | 654 (80.34%) | 722 (80.76%) |
| $S_{14\text{-}B}$ | 0 (0.0%) | 1 (0.4%) | 1 (0.11%) | 1 (0.12%) | 0 (0.00%) |
| $S_{14\text{-}C}$ | 0 (0.0%) | 1 (0.4%) | 0 (0.00%) | 0 (0.00%) | 0 (0.00%) |
| $S_{14\text{-}D}$ ($S_G$) | 1 (0.7%) | 0 (0.0%) | 2 (0.21%) | 0 (0.00%) | 1 (0.11%) |
| $L_{16\text{-}A}$ ($L_A$) | 72 (48.6%) | 18 (6.9%) | 76 (8.12%) | 61 (7.49%) | 59 (6.60%) |
| $L_{16\text{-}B}$ | 0 (0.0%) | 3 (1.1%) | 7 (0.75%) | 2 (0.25%) | 5 (0.56%) |
| **$L_{16\text{-}C}$** | **0 (0.0%)** | **13 (5.0%)** | **60 (6.41%)** | **32 (3.93%)** | **32 (3.58%)** |
| $L_{16\text{-}D}$ ($L_G$) | 7 (4.7%) | 11 (4.2%) | 65 (6.94%) | 54 (6.63%) | 60 (6.71%) |
| $L_{16\text{-}E}$ | 1 (0.7%) | 0 (0.0%) | 0 (0.00%) | 0 (0.00%) | 0 (0.00%) |
| $L_{16\text{-}F}$ | 1 (0.7%) | 0 (0.0%) | 0 (0.00%) | 0 (0.00%) | 0 (0.00%) |
| Others | 0 (0.0%) | 8 (3.1%) | 23 (2.46%) | 10 (1.23%) | 15 (1.67%) |

AF: allele frequency, CT: control, SZ: schizophrenia, BD: bipolar disorder, 5-HTTLPR: serotonin transporter-linked polymorphic region. Asian-specific L allele (L16-C) is highlighted in bold.

**Supplementary Table S7.** Case control DNA methylation analysis in BD considering the promoter activity of 5-HTTLPR

| Allele ($S_A$ background) | Diagnosis | N | CpG 3 | |
|---|---|---|---|---|
| | | | DNA methylation level (%) (mean±SD) | *P*-value (Cohen's d) |
| $S_A$ | CT | 157 | 29.6±4.9 | 0.062 (0.208) |
| | BD | 126 | 30.6±4.6 | |
| $L_A$ | CT | 34 | 30.9±4.5 | 0.860 (0.015) |
| | BD | 16 | 30.9±4.4 | |
| $L_{16\text{-}C}$/$L_G$ | CT | 52 | 29.0±4.7 | **0.005** (0.596) |
| | BD | 33 | 31.8±4.6 | |
| $S_A$/$L_{16\text{-}C}$/$L_G$ (All low-activity alleles) | CT | 209 | 29.4±4.8 | **0.002** (0.293) |
| | BD | 159 | 30.8±4.6 | |

CT: control, BD: bipolar disorder, 5-HTTLPR: serotonin transporter-linked polymorphic region. Significant *P*-values are shown in bold.

**Supplementary Table S8.** Case control DNA methylation analysis in female patients with SZ or BD considering the promoter activity of 5-HTTLPR

| Alleles ($S_A$ background) | Diagnosis | N (female) | CpG 3 DNA methylation level (%) (mean±SD) | CpG 3 *P*-value (Cohen's d) |
|---|---|---|---|---|
| $S_A$ | CT | 113 | 31.6±4.7 | 0.509 (0.126) |
| | SZ | 117 | 32.2±5.2 | |
| | CT | 106 | 35.2±4.4 | 0.531 (0.053) |
| | BD | 168 | 35.0±5.0 | |
| $L_A$ | CT | 21 | 32.6±5.2 | 0.073 (0.511) |
| | SZ | 32 | 30.0±5.0 | |
| | CT | 17 | 35.4±5.2 | 0.543 (0.199) |
| | BD | 26 | 34.4±5.0 | |
| $L_{16-C}/L_G$ | CT | 49 | 32.0±4.6 | 0.410 (0.229) |
| | SZ | 30 | 30.8±5.6 | |
| | CT | 40 | 35.6±5.7 | 0.559 (0.038) |
| | BD | 42 | 35.4±5.8 | |
| $S_A/L_{16-C}/L_G$ (All low-activity alleles) | CT | 162 | 31.7±4.7 | 0.819 (0.046) |
| | SZ | 147 | 31.9±5.3 | |
| | CT | 146 | 35.3±4.8 | 0.416 (0.055) |
| | BD | 210 | 35.1±5.2 | |

CT: control, SZ: schizophrenia, BD: bipolar disorder, 5-HTTLPR: serotonin transporter-linked polymorphic region. Significant *P*-values are shown in bold.

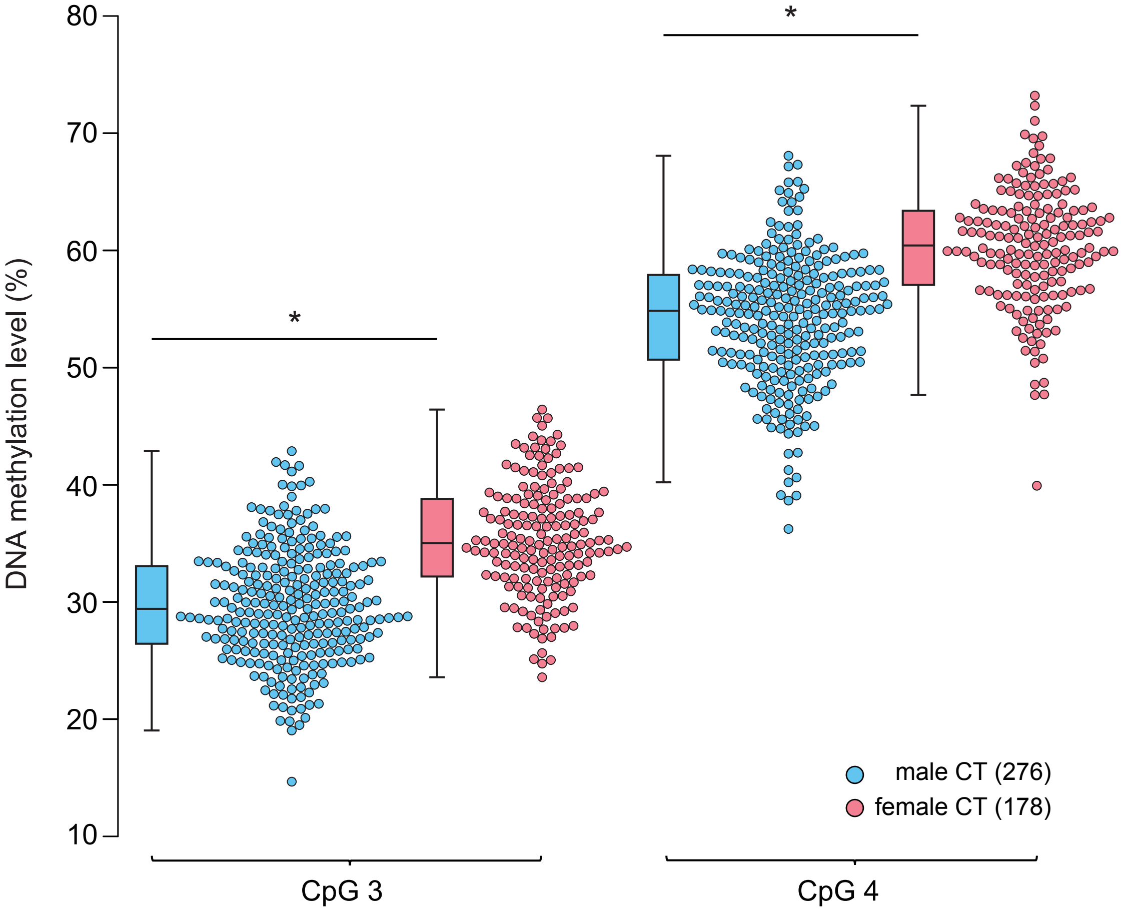


**Supplementary Figure S1. Sex-dependent DNA methylation differences in *SLC6A4*.** Comparison of DNA methylation levels between male CT and female CT. CpG3 and CpG4 showed significantly higher methylation in female CT than in male CT. $*P < 2.20E\text{-}16$.

| | | | | | | CpG3 | | | | | | | | | | | | | | | | | | | | | | | | | CpG4 | | |
|---|---|---|---|---|---|---|---|---|---|---|---|---|---|---|---|---|---|---|---|---|---|---|---|---|---|---|---|---|---|---|---|---|---|
| *Primates* | **Human** | **G** | **A** | **A** | **A** | **C** | **G** | **A** | **A** | **A** | **G** | **C** | **A** | **A** | **G** | **C** | **A** | **A** | **C** | **T** | **C** | **T** | **T** | **C** | **C** | **C** | **A** | **A** | **A** | **G** | **C** | **G** | **C** |
| | Chimp | G | A | A | A | C | G | A | A | A | G | C | A | A | G | C | A | A | C | T | C | T | T | C | C | C | A | A | A | G | C | G | C |
| | Gorilla | G | A | A | A | C | G | A | A | A | G | C | A | A | - | - | - | - | C | T | C | T | T | C | C | C | A | A | A | G | C | G | C |
| | Orangutan | G | A | A | A | C | G | A | A | A | G | C | A | A | - | - | - | - | C | T | T | T | T | C | C | C | A | A | A | G | T | G | C |
| | Gibbon | G | A | A | A | C | G | A | A | A | G | C | A | A | - | - | - | - | C | T | T | T | T | C | C | C | A | A | A | T | C | G | C |
| | Rhesus | G | A | A | A | C | G | A | A | A | G | C | A | A | - | - | - | - | C | T | T | T | T | C | C | C | A | A | A | G | C | G | C |
| | Crab-eating macaque | G | A | A | A | C | G | A | A | A | G | C | A | A | - | - | - | - | C | T | T | T | T | C | C | C | A | A | A | G | C | G | C |
| | Baboon | G | A | A | A | C | G | A | A | A | G | C | A | A | - | - | - | - | C | T | T | T | T | C | C | C | A | A | A | G | C | G | C |
| | Green monkey | G | A | A | A | C | G | A | A | A | G | C | A | A | - | - | - | - | C | T | T | T | T | C | C | C | A | A | A | G | T | G | C |
| | **Marmoset** | **G** | **A** | **A** | **A** | **C** | **G** | **A** | **A** | **A** | **G** | **C** | **A** | **A** | **-** | **-** | **-** | **-** | **C** | **T** | **C** | **T** | **T** | **C** | **C** | **C** | **A** | **A** | **A** | **G** | **A** | **G** | **G** |
| | Squirrel monkey | G | A | A | A | C | G | A | A | A | G | C | A | A | - | - | - | - | C | T | C | T | T | C | C | C | C | A | A | G | C | C | C |
| *Rodents* | Mouse | G | A | A | A | C | T | A | C | A | G | C | A | A | - | - | - | - | C | T | T | T | T | G | C | C | A | A | A | A | A | G | C |
| | Rat | G | A | A | A | C | T | A | C | A | G | C | A | A | - | - | - | - | C | T | T | T | T | G | C | C | A | A | A | A | A | A | C |

Sequence alignments of human and marmoset are shown in bold. Deviations from human sequence are highlighted in grey. Short lines indicate no base in the aligned species.

**Supplementary Figure S2. Evolutionary conservation of CpG3 in *SLC6A4*.** Sequence alignments of the genomic region around CpG3 and CpG4 in *SLC6A4*. CpG3, but not CpG4, in the *SLC6A4* is conserved among primates. Note that CpG3 is not conserved in rodents.

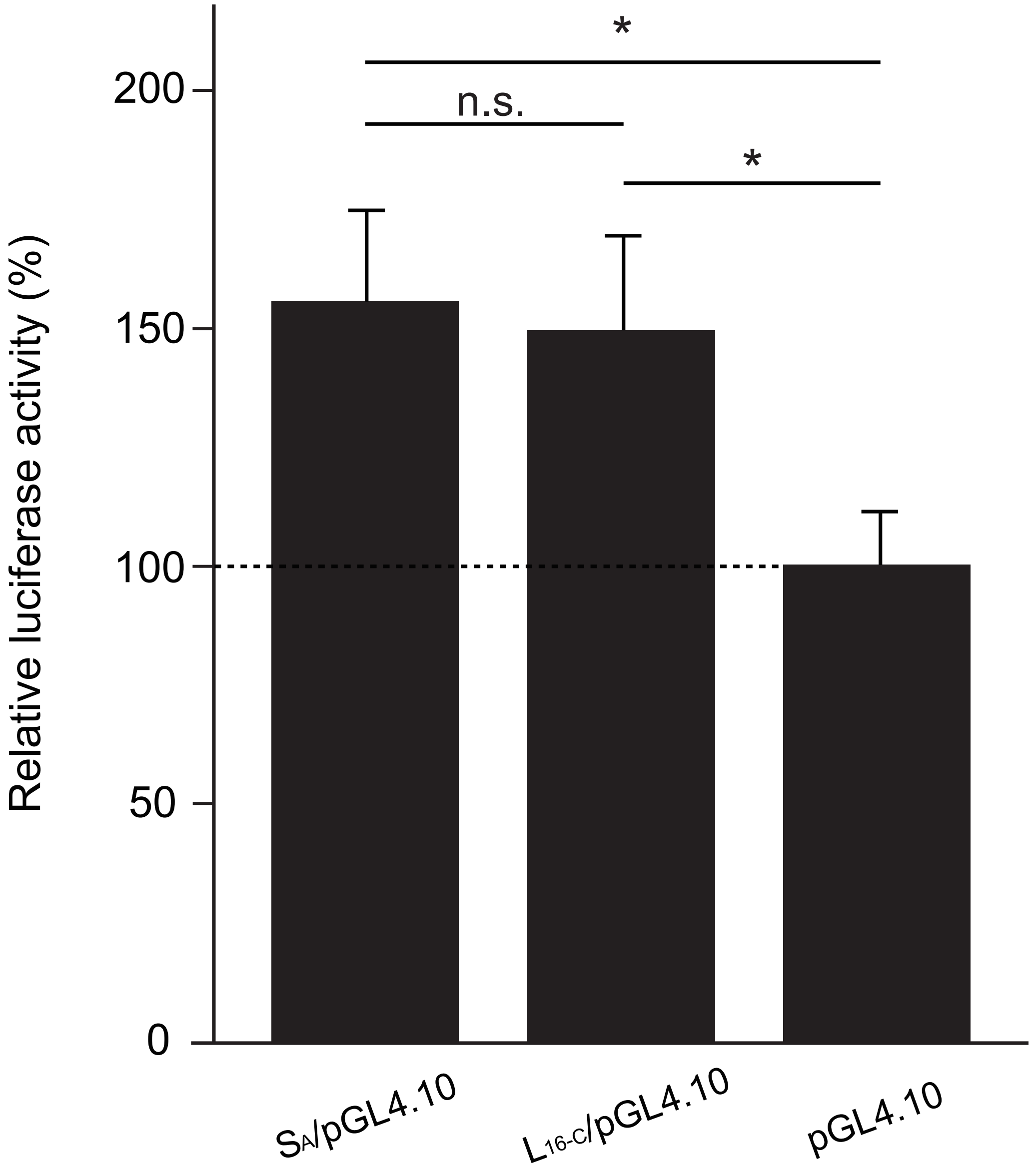


**Supplementary Figure S3. Luciferase reporter assay of $L_{16\text{-}C}$.** The effect of each allele on reporter transcription was measured as relative luciferase activity and was normalized to the mean ratio of blank control luciferase vector (pGL4.10). There was no statistically significant difference between $S_A$ and $L_{16\text{-}C}$ ($P > 0.905$, Tukey-Kramer test). *$P < 0.05$.

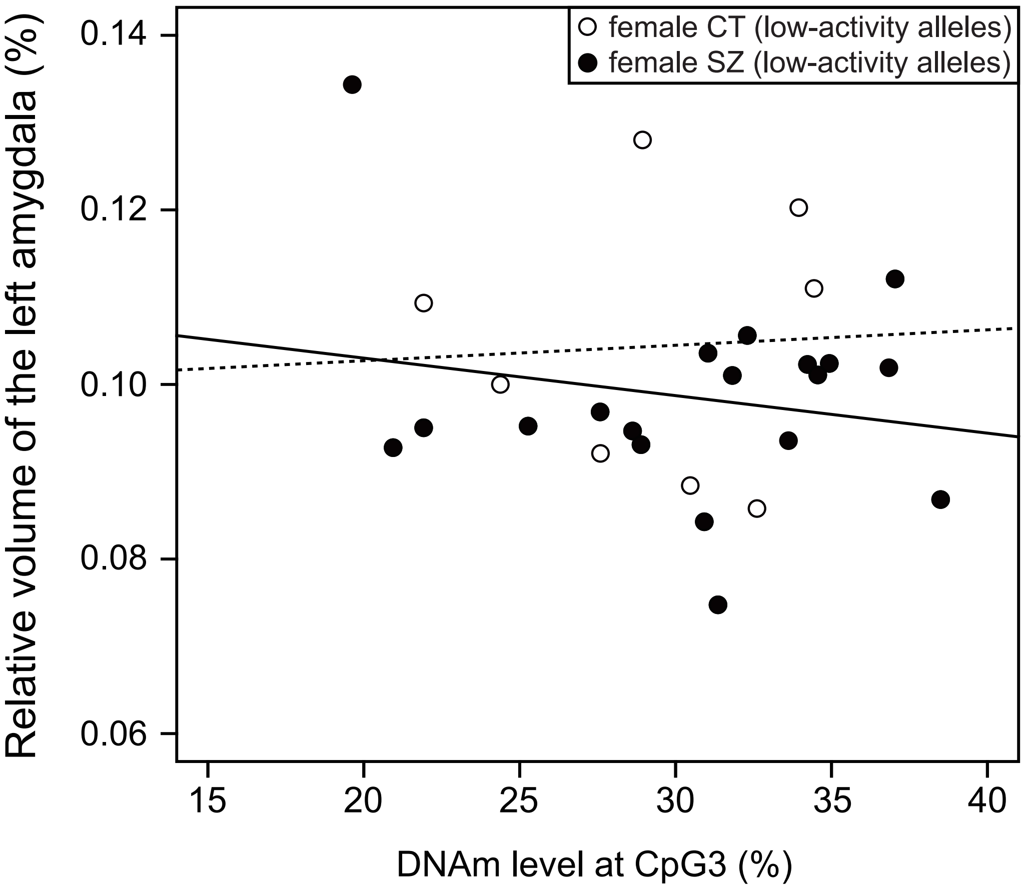


**Supplementary Figure S4. Correlation between DNA methylation levels at CpG3 and the left amygdala volume in females.** No significant correlation was observed in female patients with SZ harboring low-activity alleles (N = 19, *R* = -0.194, *P* = 0.425, solid line) or female CT (N = 8, *R* = 0.052, *P* = 0.903, broken line).
